# Proteomic characterization of major fish allergy responsive protein parvalbumins in Hilsa (*Tenualosa ilisha*): A commercially important fish in Southeast Asia

**DOI:** 10.1101/2025.06.03.657624

**Authors:** Nazma Shaheen, Zongkai Peng, Amit Singh, Asfia Wahab, Maria Gasset, Md Hafizul Islam, Oumma Halima, Aleena F Ali, Salmaan M Shah, Malek Salkini, Saaim Saleemi, Abdullah F Mallah, Ayan Khan, Sohaib Mesiya, Morshed Khandaker, Aayan Zarif, Akbar Ali, Amir Samour, John W Peters, Zhibo Yang, Nagib Ahsan

## Abstract

The protein parvalbumin (PRV)-beta (PRVB) is the primary cause of food allergies to bony fish. Although PRVB is a well-characterized protein in many bony fishes, little is known about the Hilsa, an anadromous fish with great economic importance and predominantly found in Southeast Asia. In this study, we characterized the Hilsa PRV utilizing various proteomic approaches in response to two major riverine habitats and developmental stages. Unique peptide sets corresponding to three PRV isoforms were identified in Hilsa muscle tissues. Label-free quantitative proteomic analysis coupled with ELISA revealed higher levels of PRVB in young fish compared to adults, irrespective of their riverine habitats. A comparative quantitative analysis of PRVB further demonstrated that Hilsa had less PRVB than other commonly consumed freshwater fish species. Multiple reaction monitoring (MRM)-based targeted proteomic approach showed the potential of PRV as a marker protein for allergen quantitation and authenticating the presence of Hilsa in a complex freshwater fish mixture. Our findings collectively offer fundamental knowledge on Hilsa PRVs for further investigation on the food safety and quality evaluation of Hilsa fish.

## 1. Introduction

Fish allergies, particularly those triggered by the protein parvalbumin, are a significant global health concern (Stephen et al., 2017). Parvalbumin (PRV), particularly parvalbumin-beta (PRVB) is a calcium-binding protein predominantly found in the muscle tissue of bony fish and is responsible for nearly 90-95% of fish-induced allergic reactions (Stephen et al., 2017; Wai et al., 2021; Mukherjee et al., 2023). Its unique stability with high resistance to heat, digestion, and denaturation makes it a potent allergen, capable of triggering immune responses even after fish is cooked (Tsabouri et al., 2012). Cross-reactivity among fish species is a significant concern for individuals with fish allergies, as PRVB exhibits a high degree of structural similarity across different species (Bugajska-Schretter et al., 1998; Swoboda et al., 2002; Kuehn et al., 2014, Mukherjee et al., 2021; Franciskovic et al., 2024). This cross-reactivity means that over 90% of fish-allergic individuals will likely experience allergic reactions to multiple fish species (Bugajska-Schretter et al., 1998; Swoboda et al., 2002). In contrast, cartilaginous fish, such as sharks and rays primarily express parvalbumin-alpha (PRVA), a form that is much less allergenic (Stephen et al., 2017; Mukherjee et al., 2023).

Hilsa (*Tenualosa ilisha*), the national fish of Bangladesh, is often referred to as the “king of taste” due to its rich, buttery flavor and high nutritional value, particularly its protein content and omega-3 and omega-6 fatty acids (Acharya et al., 2022). It plays a central role in the culinary traditions of Bangladesh, where it is both a dietary staple and an integral part of national celebrations. About 90% of the world’s Hilsa production comes from Bangladesh, making it the largest Hilsa-catching nation in the world (Rahman et al., 2018; Faruque and Das, 2024). The majority of the Hilsa catch (65%) comes from the marine waters of the northern bay of the Bangle, with the remaining 33% coming from the Meghna estuary system. The Padma and other inland riverine systems only comprise up 2% of the overall Hilsa production (2%) (Hossain et al., 2020). Numerous investigations revealed that Bangladesh’s main rivers, including the Padma and Meghna, have distinct physicochemical properties (Flura et al., 2016; Shaha et al., 2020; Sultana et al., 2022) that influence the growth rate and dominance of plankton populations (Hossain et al., 2020; Shaha et al., 2022), whereas estuarine phytoplankton is a significant source of food for Hilsa.

Interestingly, people of Bangladesh and West Bengal in India believe that Hilsa fish from the Padma River are tastier than those from other riverine systems. A recent study on Hilsa fish from two Indian river systems found that the better taste of Hilsa from the Padma River might be linked to higher levels of n3:n6 fatty acids and higher amounts of alanine, aspartic acid, glutamic acid, oleic acid, and palmitoleic acid (De at al., 2019).

While Hilsa is an anadromous species, it migrates between marine and freshwater environments during its life cycle (Asaduzzaman et al., 2020). This migration exposes the Hilsa fish to various environmental conditions, including variations in water salinity, diet and pollution (Shohidullah Miah, 2015; Hossain et al., 2019). According to De et al. (2019), the lipid and amino acid profiles of Hilsa fish have changed dramatically across size groups and various habitats of Indian rivers and marine environments. Additionally, there have been significant variations in the proximate composition, mineral content, total protein content, and fatty acid content of two Tenualosa sp. from three distinct east coast areas of India (Acharya et al., 2022). It has also been reported that the texture of fish muscle is strongly correlated with developmental stages and environmental factors, such as food habits (Kotzamanis et al., 2020; Leeper et al., 2022).

Regardless the economic importance and popularity, Hilsa is notable for its possible allergy risk (Das et al., 2005; Chatterjee et al., 2006). According to a recent cross-sectional self-declaration survey of 970 people, Hilsa is one of the fish species that causes the greatest allergies in Bangladesh (Khan et al., 2024). Changes in the muscle proteome of Hilsa fish in response to different riverine habitats and developmental stages are therefore not surprising, which raises the question of whether these environmental factors may influence the amounts of PRV, the primary protein that causes fish allergies (Addis et al., 2010; Mahboob et al., 2019). More importantly, to our knowledge, there is no report on the characterization of Hilsa PRV proteins. This study also aims to examine the PRV levels in young (jatka) and adult Hilsa fish in connection with the two main riverine systems of Bangladesh. Consequently, understanding the variations in PRV levels between different life stages of Hilsa fish will enhance our comprehension of their ecological dynamics. This knowledge may also inform management strategies to ensure the sustainability of Hilsa populations in the face of environmental changes and fishing pressures. Therefore, further research into the structural and functional characteristics of Hilsa PRV proteins could provide important insights into their involvement in Hilsa fish biology and potential food safety consequences. The results of our study enhanced our understanding of fish allergies and the sustainability of Bangladesh’s Hilsa fishery while also providing crucial information regarding the food safety of Hilsa fish.

## 2. Materials and Methods

### 2.1. Hilsa fish sample collection

In Bangladesh, two different sizes of live Hilsa fish were collected from two major rivers namely Meghna and Padma. The Meghna (22°50’03.5” N, 90°20’06.5” E) and Padma (24°29’32.1” N, 88°18’17.6” E) rivers are located in the south and north side of the country, respectively (**Figure 1A**).

**Figure 1.**
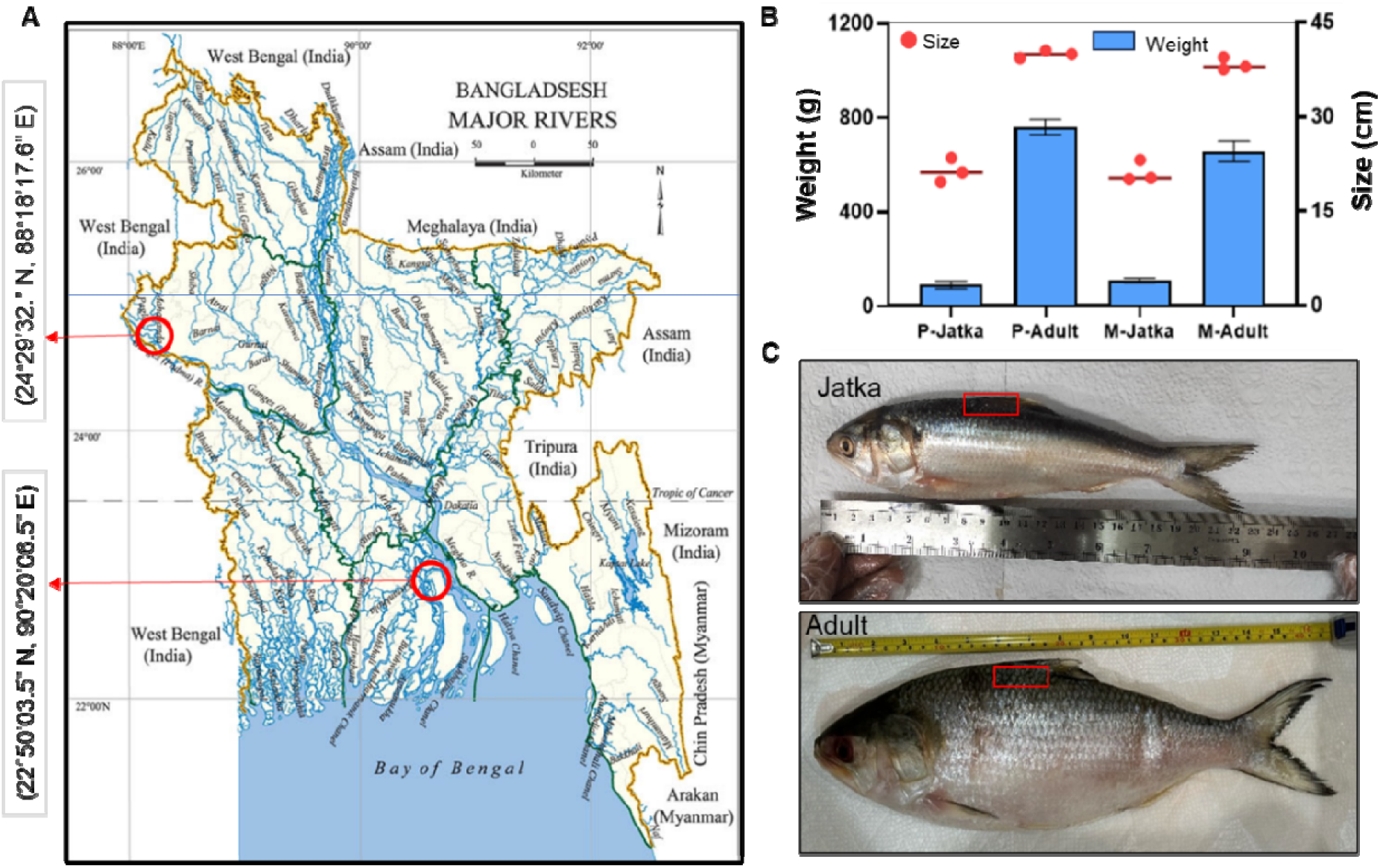
Hilsa fish collection and morphological characteristics. A, fresh Hilsa fish were caught from two main riverine system namely Meghna (22°50’03.5” N, 90°20’06.5” E) and Padma (24°29’32.1” N, 88°18’17.6” E). Red circles in the map showing the sampling sites (geographic co-ordinates and place). The map was adopted from national encyclopedia of Bangladesh (https://en.banglapedia.org/index.php/River). B, An average (n=3) size and wight of jatka (young) and adult Hilsa fish were used for proteomic analysis. P and M indicate Padma and Meghna, respectively. C, muscle tissues were collected from the dorsal portion of each fish (marked by a red box) were subjected to subsequent protein extraction and analysis.

Based on a recent study, female Hilsa from multiple populations in Bangladesh reach maturity (M50) at 31 cm total length (Abu Rayhan et al., 2023). In this study Hilsa fish having an average weight of 90.16 to 109.5 g and length of 21.2 to 21.46 cm are referred to as young (jatka) while those having an average weight of 658.26 to 761 g and length of 38.36 to 40 cm are classified as mature (adult) (Figure 1B). It has been reported that generally jatka’s are predominant in freshwater rivers, whereas adult fishes are primarily found in the estuary and marine water (Sahoo et al., 2018). For each Hilsa group three fish were selected. After collecting the live fish from the fishing boat, muscle tissue was collected from the dorsal portion of each fish (Figure 1C), as described earlier (Shaheen et al., 2023). All the required paperwork for animal ethics clearance and fieldwork was approved by the authority before the start of this study (Ref. No.: KUAEC-2021/09/20) (Shaheen et al., 2023).

### 2.2. Muscle protein extraction and proteomic analysis

Muscle proteins were extracted using RIPA (Thermo Scientific™, Cat # 89901, USA) lysis buffer and subjected to in-solution enzyme digestion according to our earlier published protocol (Shaheen et al., 2019). Briefly, an equal amount of protein (100 μg) from each sample was subjected for in-solution protein digestion using trypsin/LysC (V5071, Promega, USA) at 37°C for overnight in a rotator shaker (Cat# 13-687-717, Thermo Fisher Scientific, CA, USA). Tryptic peptides were desalted using C18 Sep-Pak plus cartridges (Waters, Milford, MA, USA) and equal amount (2 µg/sample) of digested tryptic peptides were injected for LC-MS/MS analysis. The LC-MS/MS analysis was performed using a Dionex UltiMate ® 3000 UHPLC system (Thermo Fisher Scientific, CA, USA) connected to a Q Exactive HF-X mass spectrometer (Thermo Fisher Scientific, Waltham, MA) according to our recently published protocol (Peng et all, 2024). The database search for each LC-MS/MS RAW file was performed using the Proteome Discoverer (PD) 2.4 software (Thermo Fisher Scientific, San Jose, CA) against the Hilsa protein sequence database, as was previously described (Shaheen et al., 2019). The label-free quantitative and comparative analysis of the samples was conducted using the Minora algorithm and adjoining bioinformatics tools available in Proteome Discoverer. Protein showed a 1.5-fold increase or decrease in abundance with a p-value < 0.05 was considered statistically significant.

### 2.3. Quantitation of parvalbumins using ELISA

Eleven different fish species, i.e., Atlantic salmon (*Salmo salar*), Atlantic cod (*Gadus morhua*), Mahi mahi (*Coryphaena hippurus*), Tilapia (*Pelmatolapia mariae*), Catfish (*Ictalurus punctatus*), Swai (*Pangasius bocourti*), Bighead carp (*Hypophthalmichthys nobilis*), Grass carp (*Ctenopharyngodon Idella*), and Bigmouth buffalo fish (*Ictiobus cyprinellus*) were collected from the Chinese fish market in Oklahoma City, USA. Similarly, frozen small (<500 g, n=5) and large (>800 g, n=9) Hilsa fish were collected from the local Indian grocery stores. Fish sample (1 g) was collected from the dorsal portion of each fish and stored at –80°C until further use.

PRVB protein quantitation was conducted using the Fish Parvalbumin ELISA Kit (ARG80797, Arigo, Taiwan). The recombinant mackerel (*Scomber japonicus*) parvalbumin beta (Sco j 1, P59747) (Pérez-Tavarez et al., 2021) was used as a standard to generate calibration curves. All steps followed the vendor’s protocol with minor modifications. Briefly, 100 mg of fish was homogenized in 1.5 mL of Extraction and Dilution buffer using a Bead Mill 24 (S=6.00 C=01 T=1:00, Fisherbrand, CA, USA). The mixture was then incubated at 60°C for 15 minutes, followed by centrifugation at 2000g for 10 minutes to collect the supernatant. Subsequently, a 10-fold dilution of all fish samples was performed, followed by centrifugation at 7000g for 5 minutes to obtain particle-free solutions for the ELISA experiment. Finally, the optical density (OD) was read at 450 nm (reference OD 620 nm) within 30 minutes using a microplate reader (Synergy H1, BioTek, VT), and the protein concentration was quantified based on the standard parvalbumin using a logistic curve fit.

### 2.4 Multiple reaction monitoring (MRM) analysis of Hilsa and other parvalbumin proteins

A total of seven unique peptide sequences that correspond to the PRVs of six different fish species, i.e., the Hilsa, Tilapia, Catfish, Swai, Bighead carp, and Grass carp, were selected for a targeted proteomic analysis using a triple quadrupole mass spectrometer (TSQ Quantiva, Thermo Fisher Scientific, Germany) coupled with a nano UHPLC (Dionex 3000, Thermo Fisher Scientific, Germany).

To create a complex fish mixture, 10 µg of trypsin-digested peptides from each fish species were combined in a low protein-binding Eppendorf tube. Complex tryptic peptides (2 µg) were chromatographically separated using reverse-phase chromatography with acidified ACN (acetonitrile with 0.1% formic acid) and water (0.1% formic acid in LC-MS grade water) as the mobile phases. Separation of peptides was achieved with a total 60 min run with an analytical gradient of ACN from 5% to 35% in 30 min using a flow rate of 350 nL/min through an EasySpray (3 µm, 75 µm × 15 cm, ES900, Thermo Fisher Scientific, Germany) analytical column, followed by a quick increase to 95% ACN as a wash step, then equilibrated back to 5% ACN for the remaining 10 min.

For all the peptides, the y-ions were selected for MRM analysis. The MRM method was developed using Skyline software version 24.1 (https://skyline.ms), an open-source tool for mass spectrometry analysis (**Table S1**). The parameters used for the quadrupole mass spectrometer instrument are as follows: positive ion spray voltage, 2200 kV; ion transfer tube temperature, 350 °C; cycle time, 0.5 s; chromatographic peak width, 3.0 s; collision pressure, 1.5 mTorr; and Q1 and Q3 resolutions, 0.2 and 0.7 (FWHM), respectively. MS RAW files were further processed and analyzed by Skyline software (Ver. 24.1.0.199).

### 2.5. Bioinformatics analysis

PRV sequences were downloaded from WHO/IUIS Allergen Nomenclature Database (https://www.allergen.org/pubs.php). Multiple amino acid sequence alignment was performed using an open-source sequence alignment platform Clustal Omega (https://www.ebi.ac.uk/jdispatcher/msa/clustalo) and heatmap on percent (%) identity was generated using Python. Phylogenetic analysis was performed using the neighbor-joining (NJ) method through MEGA V. 11 (Tamura et al., 2021) and visualized using iTOL V. 6 (Zhou et al., 2023). Bootstrap test percentage (≥ 60%) of 10,000 replicates (Karnaneedi et al., 2020) are shown next to the branches. Principal component analysis (PCA) and Volcano plot were generated by SRplot, a free online bioinformatic platform. Histograms were generated using python script (https://github.com/tingying-he/KaiVis/blob/main/protein-rank-bar-chart-with-highlights.ipynb). Bar plots and Violin plots were generated with GraphPad Prism 7.0.

## 3. Results

### 3.1 Identification of Parvalbumin isoforms in Hilsa muscle tissue

A total of 4,448 unique peptides corresponding to 646 unique protein groups were successfully identified and quantified from Hilsa muscle samples (**Tables S2 and S3)**. Among these, three proteins were identified as parvalbumins (PRVs) with high sequence coverage ranged between 72-74%. Each Hilsa PRV was uniquely identified by at least four distinct peptides, providing high confidence of identification of multiple PRV isoform amino acid sequences in Hilsa muscle tissue (**Figure 2A, Table S4**). The three Hilsa PRVs have extremely different N-terminal sequences, and we were able to identify these distinct peptides using our discovery proteomic approach (**Figure 2B**).

**Figure 2.**
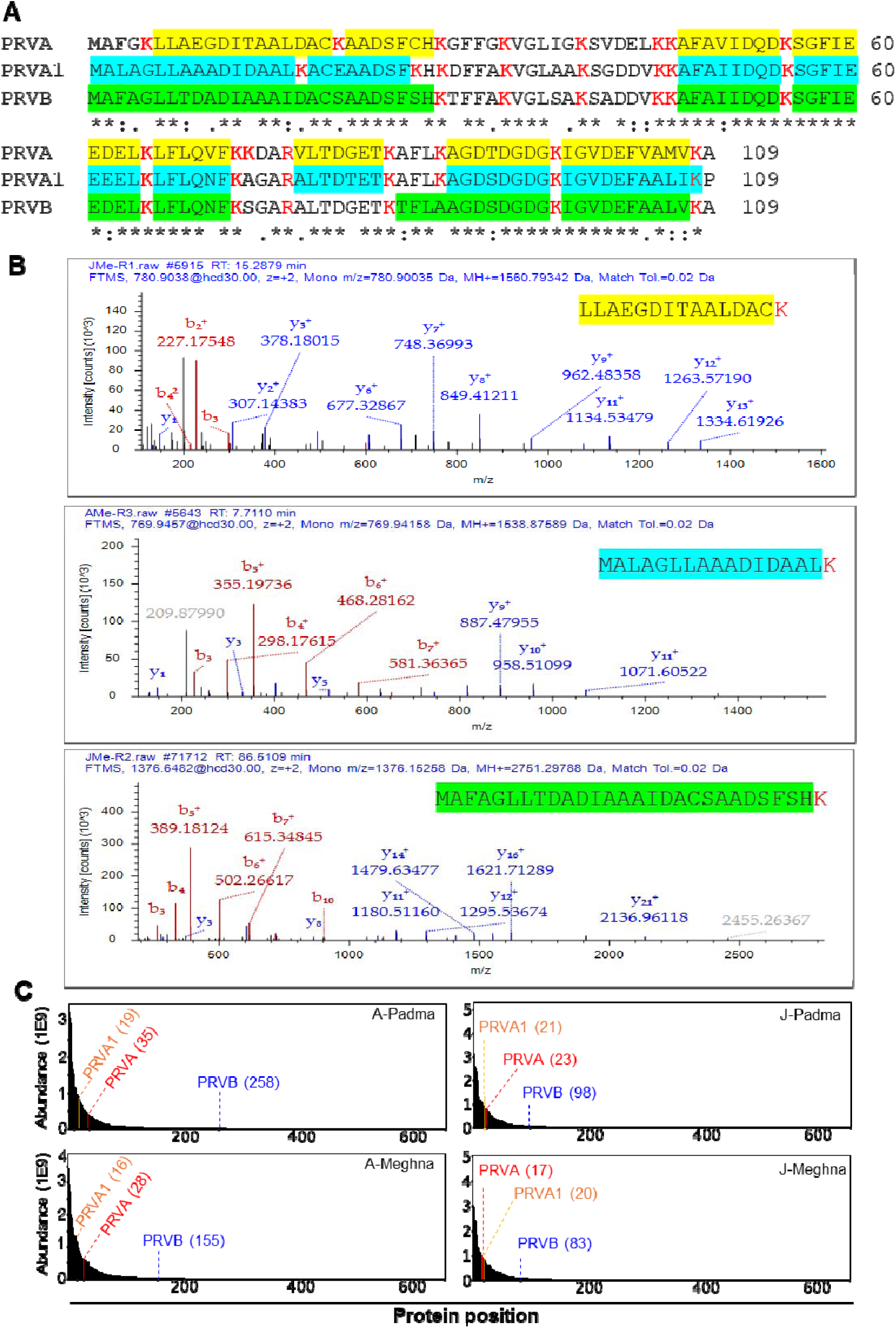
LC-MS/MS proteomic characteristics of Hilsa PRV isoforms. A, multiple amino acid sequences alignment of three PRV isoforms of Hilsa fish. Peptide sequences labeled as cyan, yellow, and green were identified by LC-MS/MS analysis. B, LC-MS/MS fragmentation three distinct peptides correspond to N-terminal of PRVA (yellow), PRVA1 (cyan), and PRVB (green) isoforms. C, histogram shows the relative abundance of Hilsa PRVA, PRVA1, and PRVB in different Hilsa samples.

Among these three PRVs, accession T_ilisha_TRINITY_DN10771_c0_g2_i1.p1 corresponding to parvalbumin beta (PRVB), whereas T_ilisha_TRINITY_DN37439_c0_g1_i3.p1 and T_ilisha_TRINITY_DN10771_c0_g1_i1.p3 both were annotated as parvalbumin alpha (PRVA). For easier naming and description, following this, we termed these three Hilsa PRVs as PRVB, PRVA, and PRVA1. An abundance-based protein quantitation analysis indicates that PRVAs are among the most prevalent comparative to PRVB in Hilsa muscle tissue, regardless of their developmental phases and geographic distribution (**Figure 2C**). Multiple sequence alignment shows that the three Hilsa PRV isoforms have 74–83% sequence identity (**Figure 3A**), despite the fact that Hilsa PRVB has a high percentage of sequences (80–89%) with other freshwater and cod PRVBs (**Figure 3A**). However, Hilsa PRVA isoforms have a higher sequence identity (>70%) with other fish species belonging to the Clupeiformes, including Pacific pilchard (Sarsa1.0101) and Atlantic herring (Cluh1.0101, Cluh1.0201, and Cluh1.0301). As expected, the known PRVAs of frog, chicken, and crocodile cluster together (**Figure 3B**), while Hilsa PRV isoforms have a lower sequence identity (<61%) with the known PRVAs (**Figure 3A**). Based on the phylogenetic and amino acid sequence analyses, Hilsa PRVB is therefore closely related to common carp (Cypc1.02), grass carp (Cteni1.0101), and striped catfish (Panh1.0101) PRVBs (**Figure 3A-B**). A side-by-side analysis of Hilsa PRVs with three well-characterized PRVA and PRVB revealed no significant differences in the theoretical pI and Mw of all three Hilsa PRV isoforms from those of other PRVs. However, the amino acid positions of the PRVB attributes are highly conserved for all three Hilsa PRVs isoforms. (**Figure 3C**).

**Figure 3.**
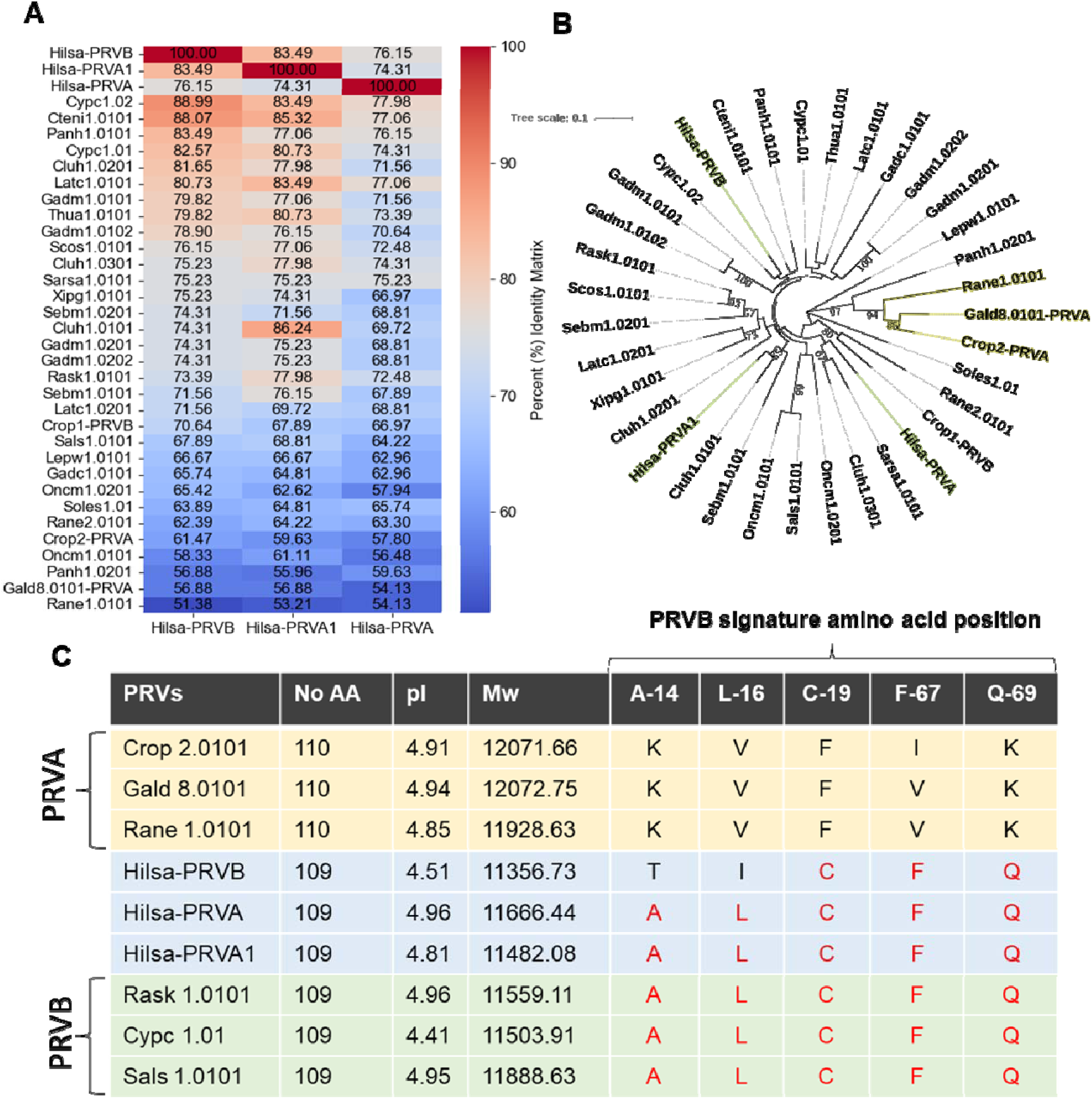
The evolutionary relationship between Hilsa PRVs and all WHO/IUIS-registered PRVs. A, heat map shows the amino acid sequence identities (%) of all WHO/IUIS-registered parvalbumins and Hilsa PRVs. The sequence identities were calculated using Multiple Sequence Alignment in Clustal Omega (EMBL-EBI). B, the phylogenetic analysis shows the relationship of Hilsa PRVs with known allergen PRVs. Hilsa PRVs are indicated by green boxes, while the yellow box shows the close clustering of chicken, crocodile, and frog PRVAs. All allergen sequences were downloaded from the WHO/IUIS website. C, comparative table shows the theoretical pI/Mw and the signature amino acid position of Hilsa PRVs compared to three well-characterized PRVB and PRVA.

### 3.2 Comparative proteome analysis of adult and jatka Hilsa collected from Padma and Meghna rivers

The label-free comparative proteome analysis of the total Hilsa muscle protein abundance shows tight clustering within the two adults (APa and AMe) and two jatka groups (JPa and JMe). However, there is a significant difference between the two sizes (adult and jatka) (**Figure 4A**). Similarly, a heat map analysis of the total number of unique proteins further demonstrates the differential abundance between the size groups (**Figure 4B**). Regardless of the two distinct riverine systems, the volcano plot analysis also showed that only PRVB, out of the three PRVs found in this study, demonstrated significant alteration (at least 1.5-fold with a P-value>0.05) between the adult and jatka fish (**Figure 4C-D**). The bar diagrams show the overall abundance of three PRVs compared to the Hilsa samples (**Figure 4E-G**).

**Figure 4.**
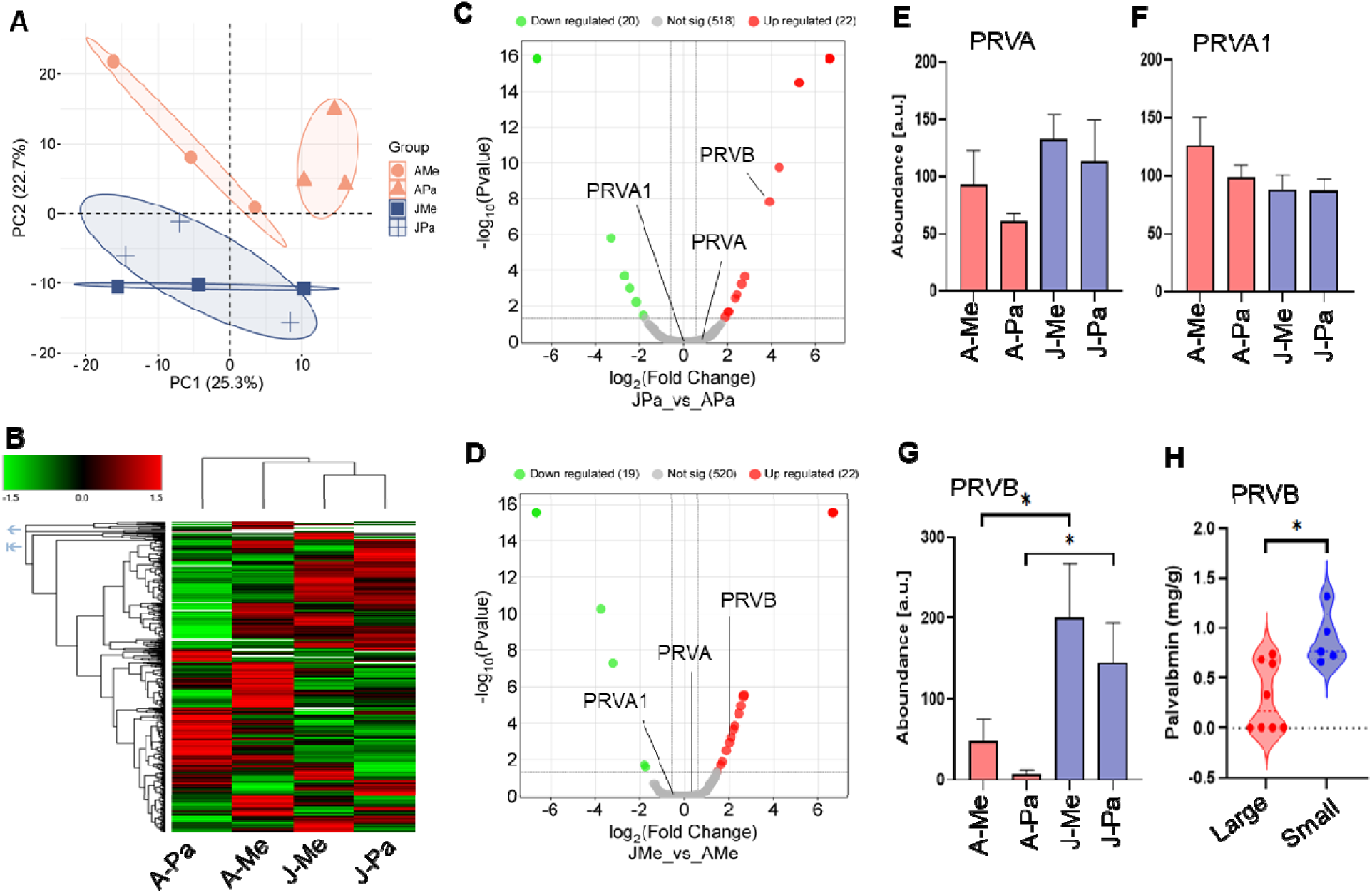
Comparative proteome profiles of adult and jatka Hilsa fish from the Meghna and Padma rivers. A, principal component analysis (PCA) of normalized total protein abundance (peak area) of adult (A) and jatka (J) Hilsa fish muscle samples collected from the Meghna (Me) and Padma (Pa) rivers. The PCA plot clearly separates the adult (AMe and APa) and jatka groups (JMe and JPa). B, Heatmap illustrates the differential expression of grouped abundance 646 proteins identified and quantified across the four groups: AMe, APa, JMe, and JPa. The color gradient reflects the relative increase (red) or decrease (green) in protein abundance, highlighting significant distinctions between adult and jatka groups. C-D, Volcano plots showing the differential protein expression between JPa vs APa (C) and JMe vs AMe (D) groups with significantly (at least 1.5-fold and adjusted p-value >0.05) upregulated (red) and downregulated (green) proteins. Gray dots represent proteins not statistically significant between the groups. E-G, bar diagrams represent PRVA (E), PRVA1 (F), and PRVB protein abundance in four Hilsa samples quantified by label-free proteomic analysis. Asterisk over the bars in PRVB indicating statistical significance between the size (adult and jatka) of Hilsa fish samples (G). H, violin plot shows the PRVB contents measured by ELISA in adult (n-9) and young (n=5) Hilsa muscle tissues.

Although there was no discernible difference in PRVs between the two riverine systems, our proteomic study demonstrated significant differences in PRVB abundance between the adult and jatka Hilsa groups. Therefore, we further intend to cross-validate the proteomic results by measuring the PRVB content in Hilsa fish samples using an independent sandwich ELISA assay utilizing an unbiased Hilsa samples pool collected from the local fish market. ELISA analysis revealed that smaller Hilsa fish with an average weight of <500 g have 1.32 mg/g PRVB, while larger Hilsa fish with an average weight >800 g have 0.62 mg/g (**Figure 4H**).

### 3.3 Comparative quantitation of parvalbumin beta in Hilsa and other fish species

To better understand the PRVB content of Hilsa fish compared to other commonly consumed freshwater and saltwater fish species, we measured the PRVB of 10 fish species using the ELISA assay. As anticipated, fish-to-fish variation in PRVB content was substantial (**Figure 5**). Compared to saltwater fish species, freshwater fish had much greater levels of PRVB. While the PRVB contents of three marine fish species (Atlantic salmon, Atlantic cod, and Mahi mahi) were extremely low (0.08-2.11 mg/g), PRVB contents were much higher (4.65-21.64 mg/g) in seven freshwater fish species (Catfish, Swai, Bighead carp, Grass carp, and Buffalo fish). Interestingly, PRVB content was extremely low in the estuarine fish Hilsa and the most cultured fish Tilapia, at 0.54 and 0.42 mg/g, respectively (**Figure 5**). Overall, except tilapia, the PRVB concentration of Hilsa fish was 40 times lower than that of other highly consumed Asian freshwater fish species.

**Figure 5.**
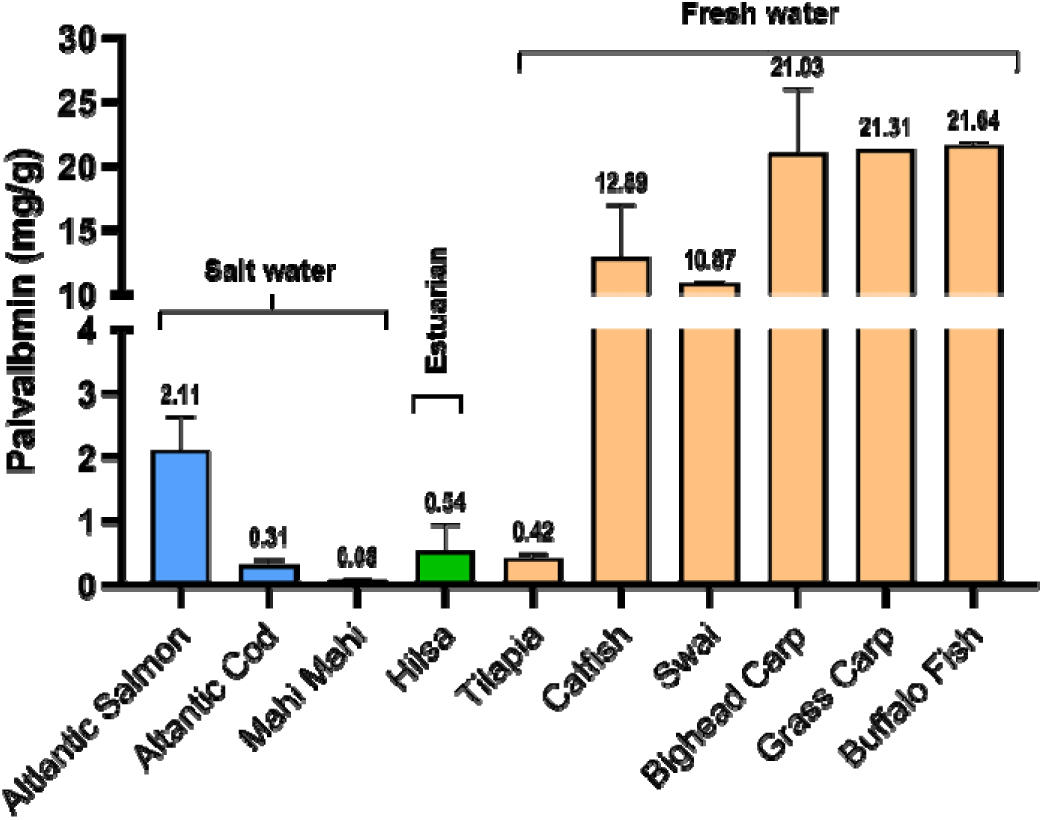
PRVB contents in the dorsal muscle tissues from 10 species of fish. Three biological samples from each of the other fish species and fourteen Hilsa fish ranging between 500 and 1000 g were used for the ELISA analysis. Each bar represents as average (±SD) and the actual value indicated above the bars.

### 3.4 Hilsa parvalbumin for targeted analysis of food safety and authenticity

A targeted proteomics experiment utilizing Multiple Reaction Monitoring (MRM) was conducted to identify the unique peptides that correlate to PRVs in six frequently ingested freshwater fish species: the swai, grass carp, Hilsa, catfish, tilapia, and bighead carp. The Hilsa peptides IGVDEFAALVK and IGVDEFVAMVK, which correspond to PRVB and PRVA, respectively, were successfully identified by our discovery proteomics approach (**Table S4**).

Unique PRV peptide sequences for all other fish species were identified using Skyline software. Except for the Hilsa PRVB peptide KIGVDEFAALVK (Yellow, **Figure 6A**), shared with tilapia, the PRV peptide sequences unique to each fish species are highlighted in green (**Figure 6A**). A trypsin-digested peptide mixture of six fish species was analyzed by LC-MS/MS using a triple quadrupole mass spectrometer. Results reveal distinct chromatographic separation with unique retention times of each PRV peptide, confirming the specificity of specific fish species (**Figure 6B**). Furthermore, the co-elution of multiple transition ions under the same retention time for each PRV peptide confirms these peptides’ optimal selection and reliability for future development of MRM-based techniques to identify and quantify PRV isoforms from any complex fish matrix (**Figure 6C**).

**Figure 6.**
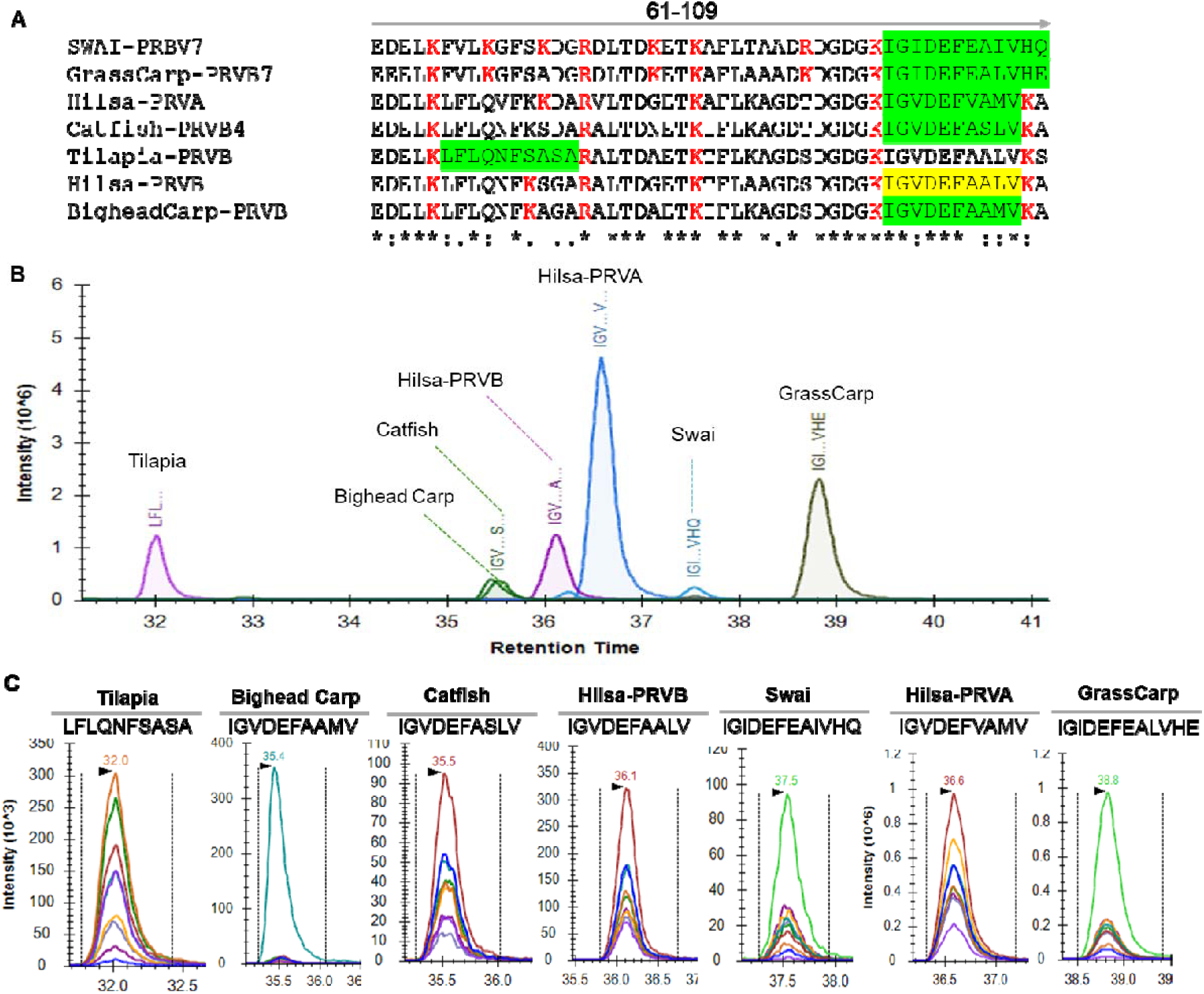
Selection and identification of ideal unique PRV peptides of six different fish species using multiple reaction monitoring (MRM) for targeted proteomics analysis. A, multiple amino acid sequence analysis of Tilapia, Bighead Carp, Catfish, Swai, Grass carp and Hilsa shows the unique (green) and shared peptides in the C-terminal of PRVs. B, chromatogram representing the seven peptides corresponding to PRVs of six different fish species. C, transition peaks on Skyline for precursor peptides LFLQNFSASA, IGVDEFAAMV, IGVDEFASLVKA, IGIDEFEAALV, IGVDEFVAMV, IGIDEFEAIVHQ, and IGIDEFEALVHE. Each color represents a different transition y ion (i.e., precursor peptide/fragment ion pair).

## 4. Discussion

Fish is one of the “big nine” sources of food allergies. Since PRVs are the primary muscle protein that causes allergic responses, risking food safety and human health worldwide. Therefore, identifying, characterizing, and quantifying the PRVs proteins in any species of fish is the first step in studying food safety and fish allergies (Ruethers et al., 2018). Like many other fish species, Hilsa has also been shown to have IgE-mediated hypersensitivity reactions (Das et al., 2005; Chatterjee et al., 2006).

More importantly, a recent survey on food allergy sensitivities found that Hilsa is one of the top three foods that cause allergic responses in Bangladeshi population (Khan et al., 2024) and that over 30% of Northern Indians patients age group 5 to 40 years experienced allergic reaction to Hilsa fish (Mandal et al., 2009).

These prior studies unequivocally demonstrated that Hilsa are allergic, just like many other bony fish species (Kobayashi et al., 2016a; Ruethers et al., 2017; Ruethers et al., 2018; Ruethers et al., 2019; Ruethers et al., 2020). However, there are currently no comprehensive reports on Hilsa parvalbumins, the primary known cause of fish allergies. Thus, a deeper comprehension of the protein sequence, isoforms, and comparative quantitative analysis of Hilsa parvalbumins in muscle tissue may result in better diagnostic tools and possible therapeutic interventions. Furthermore, this knowledge may help develop guidelines for safe consumption and management of Hilsa-related allergies among affected populations.

LC-MS/MS based proteomic approach is one of the advanced analytical platforms commonly used for identification and characterization of fish allergens (Piovesana et al., 2016; Carrera et al., 2019; Carrera et al., 2020; Carrera and Magadán, 2022; Xu et al., 2023; Liu et al., 2024). Thus far, only two proteomics studies on Hilsa fish that have been published focus on how muscle tissue (Shaheen et al., 2023) and serum (Chakraborty et al., 2024) respond to different developmental stages. Additionally, a metabolomic profile of Hilsa fish revealed that the nutritional composition of the fish is significantly altered by fecundity, geographic distribution, and growth and developmental phases (De et al., 2019). However, to our knowledge, no study has investigated how allergen proteins respond to the geographical location and developmental stage of Hilsa fish. In this work, we use LC-MS/MS techniques to characterize PRVs, the primary allergen of Hilsa fish, in relation to the geographic distribution and developmental stage.

For the first time, in this report, we revealed the actual peptide sequences of three PRV isoforms found in Hilsa muscle tissue using the discovery proteomic approach. Multiple amino acid sequence analysis coupled with phylogenetic analysis of Hilsa PRVs with WHO/IUIS-recognized PRV sequences showed that Hilsa PRVB is closely related to known freshwater fish PRVBs such as Common carp, Grass carp and Catfish PRVBs (**Figure 3A**). Interestingly, among the three Hilsa PRVs, the sequence identity was between 72-74%. It has also been discovered that several fish species, such as rainbow trout and barramundi, have intra-species lower sequence identity (<68%) of PRVs, indicating the diversity of allergenic parvalbumins (Ruethers et al, 2018). On the other hand, Hilsa PRVA and PRVA1 showed lower sequence identity (<61%) with well-characterized PRVAs of chicken, frog, and crocodile, whereas they were closely related (>75% sequence identity) to other fish PRVBs (**Figure 3A-B**). Our results agree with earlier research where PRVA and PRVB share less than 60% of their amino acid sequence (Ruethers et al, 2018). Furthermore, PRVB also differs from PRVA in terms of size, isoelectric point, and the presence of certain distinctive amino acids at particular locations (Pechére et al., 1971; Goodman and Pechére, 1977; Arif, 2009; Beale et al., 2009; Sharp and Lopata, 2014; Mukherjee et al., 2024). Additionally, our findings demonstrated that the amino acid positions of PRVB characteristics (Beale et al., 2009) are primarily conserved across all three Hilsa PRV isoforms (**Figure 3C**), suggesting that PRVA1 and Hilsa PRVA may be PRVB isoforms instead of PRVA. This conservation implies a potential evolutionary relationship among these isoforms, which may play a crucial role in the functional dynamics of the Hilsa fish. Further studies are necessary to elucidate the specific biological implications of these conserved amino acid positions in the context of PRV’s allergenicity.

Measurement of the PRV level is the first step in identifying the allergy sensitivity of a fish species (Ruethers et al., 2018). Using label-free quantitative proteomic analysis, we demonstrated that, despite the two different riverine habitats, relative PRVB abundance was significantly lower in adult Hilsa fish than in juvenile fish, whereas PRVA levels remained constant (**Figure 4C-G**). ELISA was used to further validate this result with unbiased Hilsa fish samples (both small and large) that were collected from the local market (Figure 4H). Our results suggest that the riverine environment of Bangladesh has minimal impact on PRVB, and developmental stages may regulate the Hilsa PRVB. These findings are consistent with previous studies of the protein and lipid profiles of Hilsa fish (De et al, 2019; Shaeen et al., 2023; Chakraborty et al., 2024). Due to their more significant movement activity and less dark muscle in the dorsal muscle region, smaller Hilsa fish may have higher levels of PRVB than larger fish (Kobayashi et al., 2006a; Lee et al., 2012; Kuehn et al., 2014). Future research on tissue-specific expression could further elucidate the mechanisms behind these variations and their implications for Hilsa fish allergy management.

A common phenomenon strongly linked to the allergic reaction to fish is the variation of PRVB levels in various fish species (Kobayashi et al., 2016a; Kobayashi et al., 2016b; Wai et al., 2024; Kuehn et al., 2010; Ruethers et al., 2018; Tsai et al., 2023). However, the profiling allergen of the great majority (>32K) of fish species have never been reported (Sharp and Lopata, 2014). This gap in research highlights the need for comprehensive studies that can identify and characterize these allergens. By increasing our understanding of the allergenic profiles of various fish species, we can better inform public health policies and guide individuals with fish allergies in making safer dietary choices. In this regard, we also compare the levels of PRVB in Hilsa fish with those in other commonly consumed freshwater and multiple known marine fish species (**Figure 5**). Our findings showed that PRVBs level in Hilsa fish is more than 40 times lower than that of fish species in the carp family (Common carp, Buffalo fish, and Grass carp) and more than 20 times lower than catfish-like fishes (Catfish, Swai). However, muscle tissue Hilsa PRVB levels (mg/g) are comparable to those of well-known marine fish species such as Cod, Salmon, and Mahi Mahi. Our findings are consistent with earlier studies showing that freshwater fishes had higher levels of PRVB than marine or migratory fish species. (Let et al., 2012). It has been reported that patients with parvalbumin-specific IgE alone often tolerate low PRVB fish, such as swordfish or tuna (Kuehn et al., 2010; Griesmeier et al., 2010). Therefore, the discovering various low-parvalbumin fish species could offer new dietary choices for individuals with moderate to high threshold dose reactivity (Klueber et al., 2019).

PRV corresponding unique peptides have been effectively utilized as distinctive markers of fish allergen to species-specific due to their high sequence diversity (Carrera et al., 2006; Minkiewicz et al., 2012; Carrera et al., 2013; Bucholska and Minkiewicz, 2016). In addition to identifying major fish allergens and verifying fish products, a single peptide marker can be used to develop alternative LC-MS-based assays, such as AQUA (Absolute Quantification) MRM technology, to measure allergen levels in a complex combination (Ahsan et al., 2016; Carrera et al., 2020; López-Pedrouso et al., 2023; Perkons et al., 2023). Here, using a targeted proteomic approach, two distinct peptides that correspond to Hilsa PRVs were further validated as potential Hilsa markers from a freshwater fish mixture comprising Grass carp, Bighead carp, Swai, Catfish, Tilapia, and Hilsa meat (**Figure 6**). This targeted proteome analysis not only advances our understanding of the unique PRV peptide patterns associated with each fish species, but it is also a crucial step in the identification and differentiation of fish products for a variety of commercial and culinary applications. These findings further highlight the potential of LC-MS/MS as a trustworthy instrument for assessing the quality and authenticity of fish in the seafood sector.

## 5. Conclusion

In this study for the first time, we define the PRVs of the Hilsa fish, the most economically important fish species in South Asian countries, particularly Bangladesh. We discovered three distinct PRVs isoforms in Hilsa muscle tissue using LC-MS/MS-based proteomics analysis. Based on evolutionary analysis, Hilsa PRVB is closely associated with freshwater fish PRVBs such as grass carp and common carp.

Quantitative analysis indicates that smaller fish have higher PRVB than larger Hilsa fish. Comparative analysis revealed that Hilsa’s PRVB level is 20–40 times lower than other common freshwater fish species. The discovery of distinct PRV peptides further demonstrates the versatility of the LC-MS/MS proteomics technique in characterizing allergen proteins in fish species. Unique Hilsa PRV peptides not only provide a dependable platform for detecting and quantifying the Hilsa samples in processed fish products, but also offer great potential for assessing food safety and lowering the risks of allergen contamination in packaged fish consumed globally. Since Hilsa has a lower PRVB than other popular freshwater fish species, incorporating Hilsa fish in diets may offer a sustainable way for individuals with PRVB-sensitive allergies to reduce allergy-related risks. Furthermore, the management of Hilsa fisheries will significantly benefit from the comparative quantification of PRVB levels in Hilsa fish to select the right size of fish.

## Author contributions

Conceptualization, N.A, N.S; sample collection and processing, N.S, M.H.I, O.H, M.K, J.W.P, proteomics analysis and bioinformatics, N.A, Z.P, A.W, A.S, Z.Y; protein sample preparation and ELISA, Z.P, A.F.A, S.M.S, M.S, S.S, A.F.M, A.K, M.G, S.M, A.Z, A.A, A.S; primary draft of the manuscript, N. A; N.S; writing, review, and editing, N.A, N.S, M.G, J.W.P, Z.Y; All authors have read and agreed to the published version of the manuscript.

## Data Availability

All LC-MS/MS RAW files and results files can be found at the MassIVE database (https://massive.ucsd.edu) under the following accession MSV000096794.

## Acknowledgments

N.A. gratefully acknowledges the initial funding support from the OU VPRP Office for establishing the Proteomics Core Facility. N.S acknowledge the Centennial Research Grant (Reg/Admin-3/47979) provided by the University of Dhaka to support this study. M. G acknowledge the Grant CPP2023-010544 funded by MCIN/AEI/10.13039/501100011033 and, as appropriate, by “ERDF A way of making Europe”, by the “European Union” or by the “European Union Next Generation EU/PRTR”. J.W.P. is supported by the National Institutes of Health (1R01GM138592-05).

## Supporting Information

**Table S1**. Detailed characteristics of parvalbumin peptides used for MRM analysis.

**Table S2**. List of peptides identified by LC-MS/MS analysis.

**Table S3**. List of Hilsa proteins identified by using LC-MS/MS analysis.

**Table S4**. Parvalbumin peptide sequences of hilsa fish identified by LC-MS/MS analysis.

